# DNA methylation meta-analysis confirms the division of the genome into two functional groups

**DOI:** 10.1101/2022.01.10.475724

**Authors:** Lev Salnikov, Saveli Goldberg, Parvathy Sukumaran, Eugene Pinsky

## Abstract

Based on a meta-analysis of human genome methylation data, we tested a theoretical model in which aging is explained by the redistribution of limited resources in cells between two main tasks of the organism: its self-sustenance based on the function of the housekeeping gene group (HG) and functional differentiation, provided by the (IntG) integrative gene group. A meta-analysis of methylation of 100 genes, 50 in the HG group and 50 in IntG, showed significant differences (p<0.0001) between our groups in the level of absolute methylation values of genes bodies and its promoters. We showed a reliable decrease of absolute methylation values in IntG with rising age in contrast to HG, where this level remained constant. The one-sided decrease in methylation in the IntG group is indirectly confirmed by the dispersion data analysis, which also decreased in the genes of this group. The imbalance between HG and IntG in methylation levels suggests that this IntG-shift is a side effect of the ontogenesis grownup program and the main cause of aging. The theoretical model of functional genome division also suggests the leading role of slow dividing and post mitotic cells in triggering and implementing the aging process.

## Introduction

DNA methylation, which affects chromatin state and gene expression, is one of the most studied processes of genomic regulation today [1, 2]. The specifics of the regulatory function of methylation continue to be studied, but there is no doubt that this process is related to ontogenetic development and aging [3,4]. The main direction of research in this direction focuses on the relationship between methylation and biological age in *Mammalia* [5,6,7,8]. The undoubted connection between methylation and aging processes provides an opportunity for experimental verification of various approaches to this problem. Data on the correlation between the methylation profile and biological age are also widely used as an efficiency criteria in works, aimed at rejuvenation of organism cells [9,10,11]. Interest in DNA methylation has led to the emergence of a large number of studies devoted to genome methylation and the generation of accessible databases, allowing it to be used for testing different points of view on the aging processes. Currently, there are many theories that give their explanation of the causes of aging, but there is no unanimous opinion on the problem [12,13,14]. We share the viewpoint that in the case of a multicultural organism, aging is always accompanied by a decrease in the resources required by its cells for repair and tissue regeneration [15,16,17, 18]. The reasons for such a decrease in resources are well explained in our theoretical model of the functional division of the *metazoan* genome [19,20]. We argue that the genome of all cells of the organism contains two functionally independent parts. It is the most conserved part of the genome, which provides all internal needs of any cell, or housekeeping genes (*HG*). One of the main criteria of this group was the stability of the genes included in it [21,22]. We proposed to identify into a separate functional group the genes, responsible for all specialized structures and their functions, making the organism a single integrated whole (*IntG*). The main principle we propose in our theoretical model, is the evolution base property of all IntG genes, which gives them an advantage in the consumption of cellular resources. Given the limitation of intracellular resources, such an advantage always leads to a decrease in the number of proteins, required for cell repair and produced by the HG genes. This property is essentially a manifestation of the antagonistic pleiotropy, described by Williams [23]. The difference of our theoretical model is that we assign pleiotropic properties not to individual genes, but to the entire IntG *group*, functionally isolated by us. According to our hypothesis, it is the IntG advantage in competition for resources in the cell that is the primary cause of aging. The development and maturation of the organism in *metazoans* is under the control of the ontogenesis program. We are convinced that there is no *aging program*. There is a program for *growing up, and aging is a side effect of it*. The presence of such a program is reflected primarily in the stable lifespan in various *metazoan* species [24,25]. We regard the ontogenesis program as a strictly defined sequence of IntG gene expression, starting from the zygote stage, for the purpose of organism formation. The main goal of the ontogenesis program is to achieve the maximum competitive advantage by the time the organism reaches the state of fertility, since natural selection is aimed precisely at this stage of *metazoan* development [26,27,28,29,30]. Genome methylation, as a regulatory tool, is directly involved in the development and implementation of the ontogenesis program, including the processes of cell differentiation in *Mammalian* [31].

In addition, of interest to us is the degree of inheritability of gene expression regulation expressed in their variance, as has been shown previously [32]. We are also interested in what changes occur with the baseline, or inherited diversity of epigenetic gene tuning in the genome groups we study, based on methylation data.

To summarize, the use of a meta-analysis of genome methylation in *Mammalian* allows us to test our assumptions using publicly available experimental databases. In this work, we set ourselves the following goal: using mathematical statistics, to obtain results, capable of verifying our theoretical model on the basis of genome methylation data. To do this, we need to get answers to the following questions:

1. are there statistical differences in the absolute level of methylation between the HG and IntG parts of the genome that we isolated, comparing groups in terms of both the level of methylation of gene bodies and their promoters;
2. are there any significant differences in age-related dynamics of methylation levels between these parts of the genome?
3. what is the variance of the data in the highlighted groups.

## Materials and Methods

Genome methylation data was obtained from an open source EWAS data hub (https://ngdc.cncb.ac.cn/ewas/datahub). Gene methylation results from *Whole blood* data were used. This section not only presents the most averaged values, but also contains the largest amount of data for all ages. A sample of 100 genes from the database was used for the study. The genes were divided into two groups of fifty genes each. Data on the methylation of the gene *bodies* and its *promoters* were analyzed separately. The IntG group included genes, whose activity is present in post mitotic cells and cells with low MI, which form the tissues that make up the main body mass. The TiGER open database (http://bioinfo.wilmer.jhu.edu/tiger) was used to compile the gene list. Genes from the Tissue-Specific Genes based on Expressed Sequence Tags (ESTs) section of the database were selected. The genes were selected using randomly generated numbers. The following permanently active genes from the listed tissues were included in the first group:

**Bone —** IBSP, COL2A1, ITGA11, MATN3, COL5A1, KIF23.

**Kidney —** UMOD, ARSF, KLK1, SALL1, RBKS, ACE2, TRPM6, VCAM1, DZIP1,

TMEM12.

**Liver —** ALB, BAAT, FMO3, F13B, A1BG, GNMT, SPTA1, ARG1, APOA1, HBM.

**Muscle —** MYLK2, TNNI2, TNNC2, CKM, MYOT, TNNT3, TNNT1, MYH2, ACTN3.

**Brain —** MOG, GFAP, OPCML, MOBP, GALNT9, GRIN1, NGB, NLGN3.

**Heart Muscle —** ANKRD1, NPPA, MYBPC3, MYH6, NPPA-AS1, SYPL2, SMPX.

Selection into the HG gene group was performed according to the major groups of genes, listed in the literature and categorized by the authors as “ housekeeping genes” [21,22]. Genes in the first half of the list of each group, using randomly generated numbers, were included in the group. If no data were available in the EWAS database, some of the genes in both groups were replaced by the genes next in the list. The second group included the genes represented in the indicated functional groups:

**Protein processing —** CAPN1, CAPN7, NACA, NACA2.

**Translation factors –** EIF1AD, EIF1B, EIF2A.

**tRNA synthesis –** FARSB, GARS, HARS.

**Ribosomal proteins –** RPL5, RPL9, RPL10A, RPL11.

**RNA polymerase –** POLR1C, POLR1D, POLR1E, POLR2A, POLR2B.

**Histone** - HIST1H2BC, H1FX, H2AFV, H2AFFX.

**Cell cycle -** XRCC5, RAB14, RAB18, RAB1A.

**Lysosome –** CTSA, LAMP1, LAMP2.

**Regulation of Growth and Ontogenesis** – LHFPL1, LHFPL4.

**Cytoskeletal –** ANXA1, ANXA11, ARPC1A, ARPC2.

**RNA splicing** – BATF1, DDX39B, HNRPA1P2, PABPN1,SRSF3.

**DNA repair/replication -** MCM3AP, XRCC5.

**Metabolism -** PRKAG1, B3GALT6, GSK3B, TPI1, LDHA,TALDO1, TSTA3.

## Results

Data analysis was performed using SAS 9.4 (SAS Inst Inc, Cary, NC). Methylation dynamics were assessed using linear regression analysis. The T-test was used to compare the means, and the F-test was used to compare the slopes of the regression. All p-values were based on a two-tailed hypothesis test, with values less than 0.05 considered statistically significant.

Let us examine the results of the ratio of absolute values of the methylation level between gene *bodies* and their *promoters* within the groups we identified. In the HG group, the level of methylation of gene bodies was higher than that of promoters: HG (Mean) bodies(0.3560) vs. promoters was (0.2402) p<0.0001. Difference of slopes was not significant p=0.3571. In the IntG group, the level of methylation of gene bodies was also higher than that of promoters: InfG (Mean) bodies (0.6179) vs. promoters (0.5553) was p<0.0001. Difference of slopes was significant p=0.0021. Next, we present the results of comparison of the groups in terms of absolute values of methylation levels, assessing them by the values obtained from the data on gene bodies and their promoters.

The results of comparison of the groups by the level of methylation of gene *bodies* are presented in Table 1

**Tab. 1.** Absolute values of methylation levels of genes bodies.

| The whole period of life |  | IntG |  |  | HG |  | Difference between IntG and HG |
| --- | --- | --- | --- | --- | --- | --- | --- |
|  | N | Value +/-<br>Standard error | p-value | N | Value +/-<br>Standard error | p-value | p-value |
| Mean | 84985 | 0.6179+/-<br>0.000774 |  | 86643 | 0.3560+/-<br>0.000791 |  | <0.0001 |
| Slope per year |  | -.000133+/-<br>0.000030 | <.0001 |  | -.000163+/-<br>0.000031 | <.0001 | 0.4870 |

The following graphs illustrate the changes in *means* of absolute methylation values of the genes bodies in the groups IntG and HG depending on the age.

The results of comparison of the groups by the level of methylation of genes promoters are presented in Tab 2. and Fig 2.

**Tab. 2.** Absolute values of methylation levels of genes promoters.

| The whole period of life |  | IntG |  |  | HG |  | Difference between IntG and HG |
| --- | --- | --- | --- | --- | --- | --- | --- |
|  | N | Value +/- Standard error | p-value | N | Value +/- Standard error | p-value | p-value |
| Mean | 84985 | 0.5553+/- 0.00119 |  | 81249 | 0.2402+/- 0.00103 |  | <0.0001 |
| Slope per year |  | -.000301+/- 0.000047 | <.0001 |  | -.000116+/- 0.000040 | 0.0037 | 0.0026 |

The comparison of the studied groups, as shown in Table 1,2 and Fig.1,2 shows the average value of the absolute level of methylation in the IntG and HG groups differ significantly, both in the level of methylation of gene bodies and their promoters (p<0.0001). There was also a significant difference in the observed rate of methylation decrease between the IntG and HG groups (p<0.0001). Also, the level of methylation differed significantly at each metric age point. At the same time, the difference in age dynamics was more pronounced for promoter methylation, where the difference in slope over each subsequent 5 years between IntG and HG was significant (p=0.0026). Note that similar small fluctuations in the values seen in the graphs are related to the fact that we used data from the same group of subjects in the gene groups we studied. We also examined the variation of gene promoter methylation diversity within our selected groups individually and in comparison with each other. Standard variance (STD) was considered as a measure of diversity. The results are presented in Tab.3 and Fig.3

**Fig. 1.**
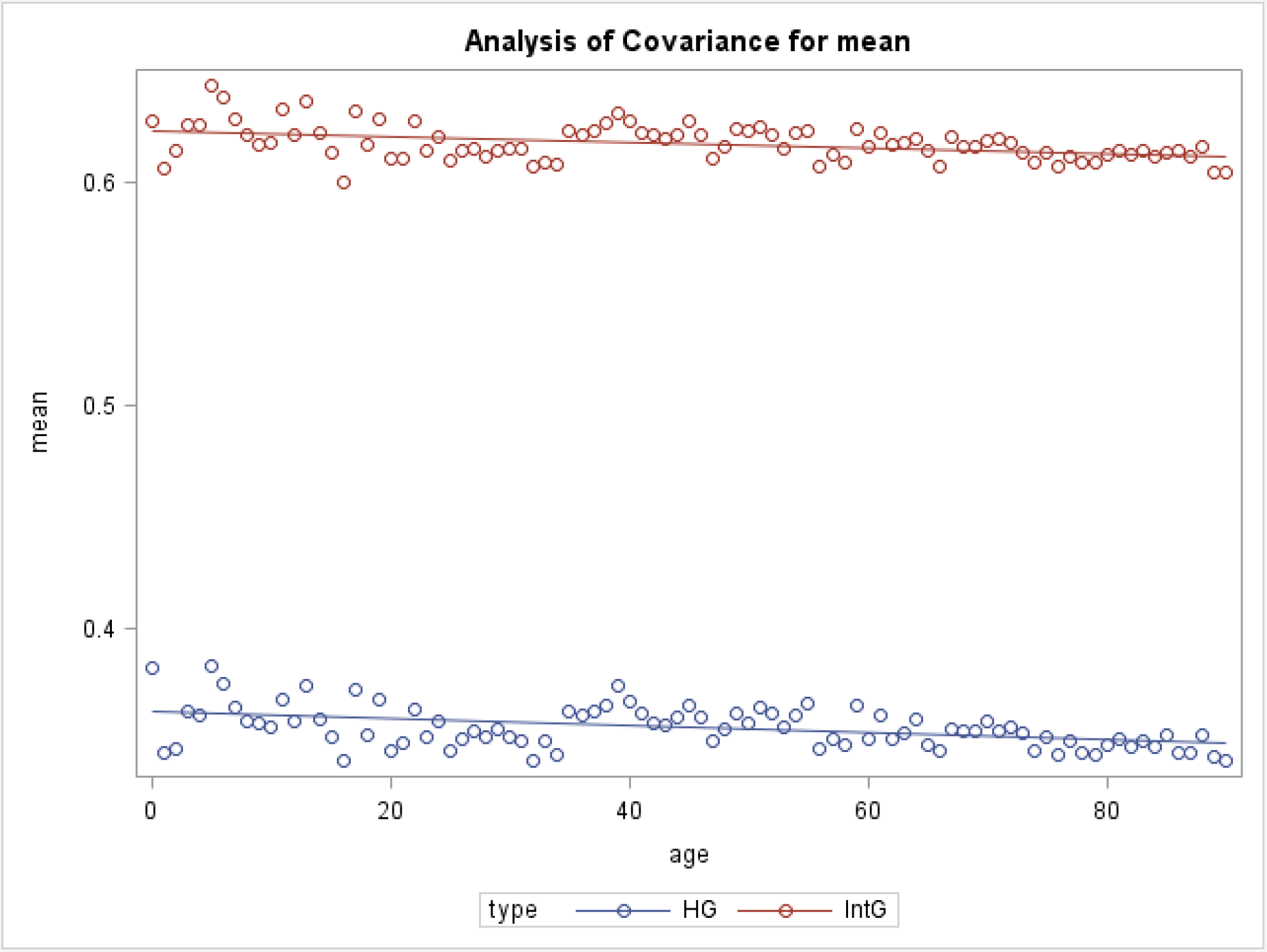
IntG vs. HG methylation of the genes bodies values per year.

**Fig. 2.**
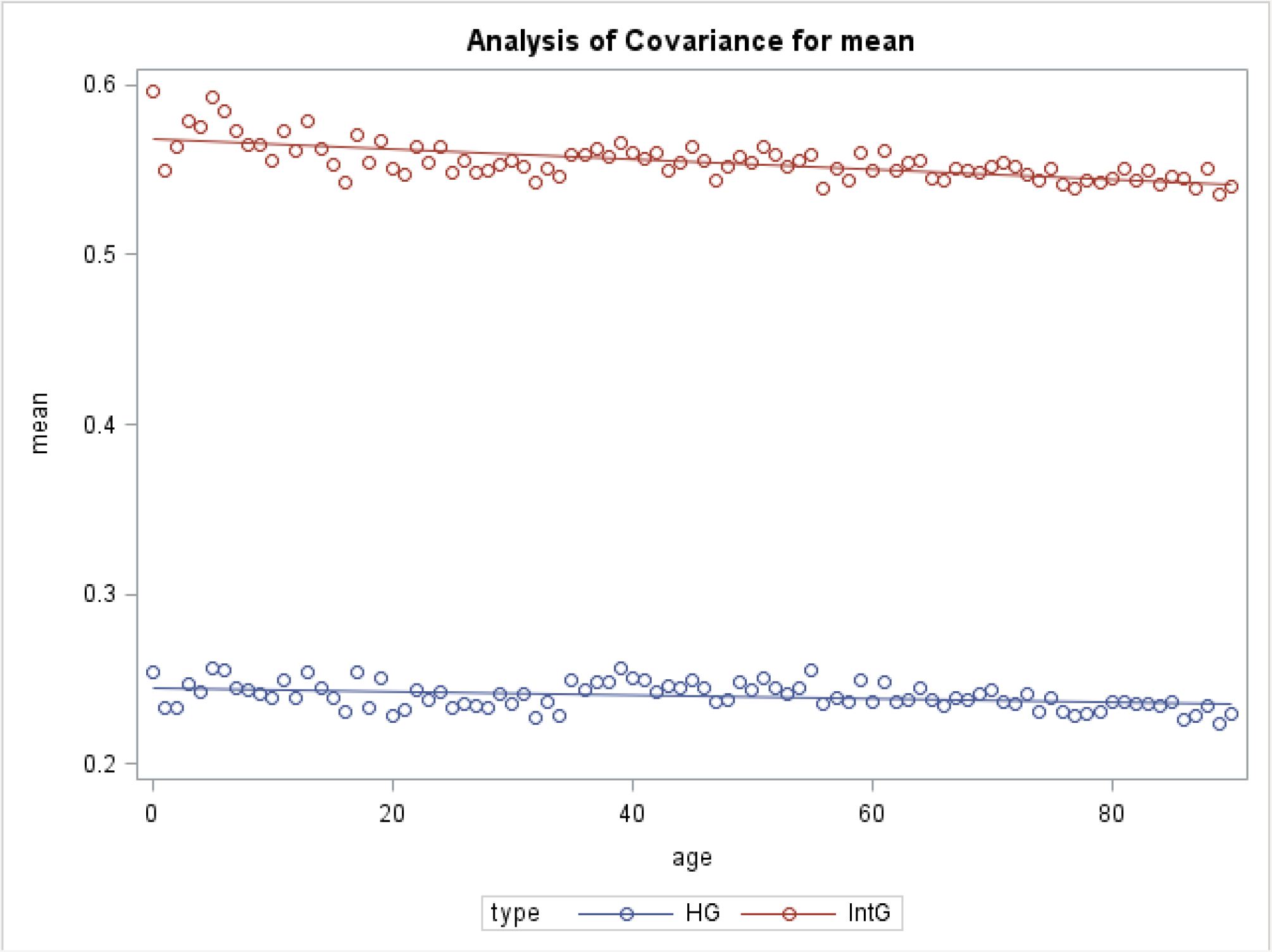
IntG vs. HG methylation of the genes promoters values per year.

**Tab3.**
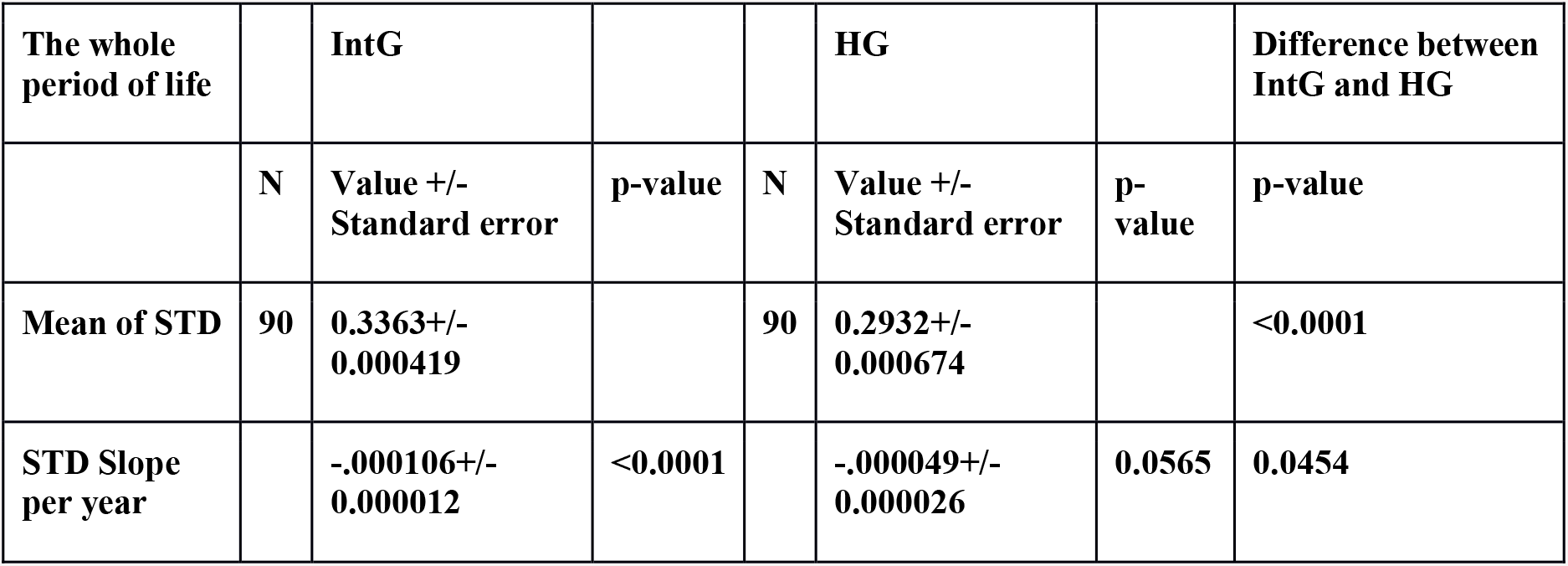
Standard deviation of methylation of promoters in HG and IntG values.

| The whole period of life |  | IntG |  |  | HG |  | Difference between IntG and HG |
| --- | --- | --- | --- | --- | --- | --- | --- |
|  | N | Value +/- Standard error | p-value | N | Value +/- Standard error | p-value | p-value |
| Mean of STD | 90 | 0.3363+/- 0.000419 |  | 90 | 0.2932+/- 0.000674 |  | <0.0001 |
| STD Slope per year |  | -.000106+/- 0.000012 | <0.0001 |  | -.000049+/- 0.000026 | 0.0565 | 0.0454 |

**Fig. 3.**
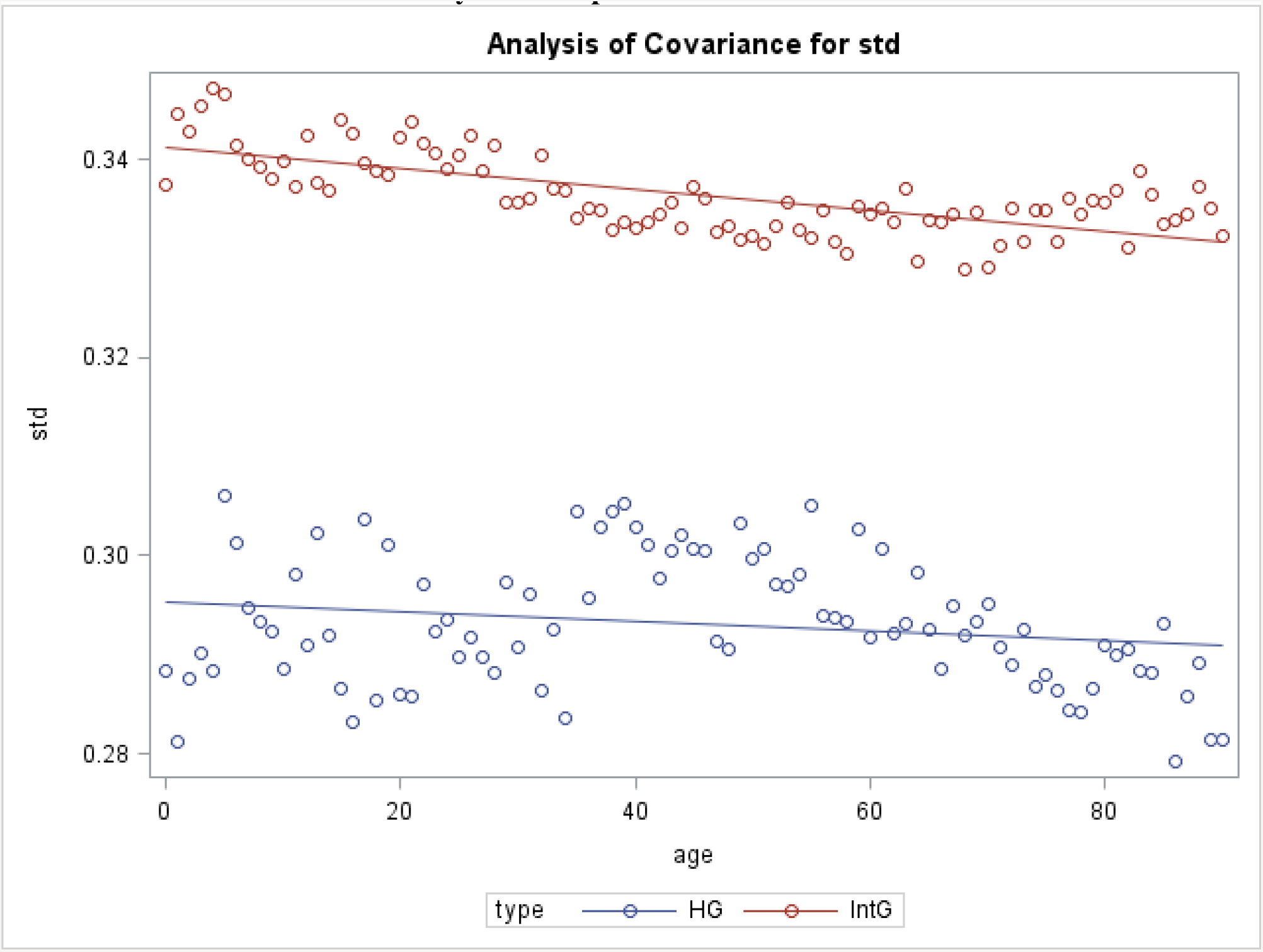
Standard deviation (STD) of methylation of promoters in HG and IntG values.

As shown in Table 3 and Fig. 3, the mean STD in the IntG group is significantly different from that in the HG group (p < 0.0001) and the STD value decreases with age (p=0.0454), repeating the trajectory of the methylation process itself. In the HG group, the variance remains practically unchanged, confirming its ontogenetic stability.

## Discussion

Discussing our results, we can conclude that our data on genome methylation supports the main statements of our theoretical aging model. For us, the approach in which we consider not individual genes but their ontogenetically determined groups is fundamental. The existence of such groups is confirmed not only by the difference between them, but also by the dynamics of their methylation with age. The data we obtained allow us to provide the following answers to the questions posed at the beginning of this work:

- statistical differences in methylation levels between our isolated HG and IntG parts of the genome are reliable for both gene bodies and their promoters throughout the entire observation period (p<0.0001);
- in the IntG gene group there was a significant decrease in methylation levels throughout the whole period of observation, especially for promoters and proved to be statistically significant (P<0.0001);
- the variance of promoter methylation data in the IntG group is statistically significantly different from HG and decreases with age, repeating the trajectory of the methylation process itself;

A certain number of explanations should be given to the presented results. As already noted, the two functional groups of genes we identified, have significant valid differences between them during the entire observation period in terms of methylation level. As the results showed, the methylation level in HG remains almost stable during the observation period. Initially, we assumed that the level of methylation in the HG genes would increase with age, but the data showed it to be unchanged with data variance remaining at the same level. In contrast, IntG methylation levels steadily decrease with age, particularly in the promoter genes. The currently available data on the relationship between methylation level and gene biosynthesis magnitude are inconsistent [33,34,35,36,37] and do not allow us to draw an unequivocal conclusion about an increase in IntG gene expression due to a decrease in their methylation level with age. When discussing the role of IntG genes, which represent the basis for organismal functions, we should mention the hyperfunction theory [38], according to which the metabolism and energy consumption of the organism must constantly increase. In contrast to this theory, for us the leading role in aging processes is played not by a direct and continuous gain of functions in the organism, but by the increase in their resource consumption, which increases after the organism reaches fertility. According to recent data [39], the level of metabolism reaches a stable value after puberty at the age of about 20 years and remains unchanged, gradually decreasing after 55-60 years when the deficit of self-sustaining functions begins to show.

By posing the question about the magnitude of the variance of methylation data in the selected groups, we wanted to find out whether there are characteristics that are unique to the IntG genes. It was found that the variance of gene promoter methylation data in the IntG group is significantly different from the HG group and decreases with age, repeating the downward trajectory of the methylation process itself. The revealed coordinated decrease in the variance of promoter methylation values with age indirectly indicates the presence of specific properties of the IntG group only.

Let’s consider the prospects and directions of our future work. In the research on aging a great attention is paid to the cells and tissues with high mitotic index (MI). This parameter allows such cells to retain a high regenerative potential and make them a target for experiments aimed at rejuvenation [40,41,42,43]. From our point of view, only influencing the epigenetic mechanisms of cells with high MI is clearly insufficient to achieve true rejuvenation of the Mammalia organism [44]. The organs and tissues consisting of postmitotic cells and cells with low MI constitute the main body mass. The survival of this group of cells is based on intracellular repair processes. For these processes, the balance of HG activity between IntG plays a crucial role. We hypothesize that the conserved advantage of IntG genes in resource consumption, shifts the balance of consumption in their direction. This is particularly significant for postmitotic cells and cells with low MI, in contrast to cells with high MI, which compensate for this problem simply by constant division. In our opinion, it is the group of cells with low MI, constituting the basis of the body mass, that plays the leading role and triggers the entire cascade of changes accompanying aging.

In conclusion, we note that our data confirm the basic statement of our theoretical model of aging - the presence of two functional parts of the genome with different behavior during a lifetime. At this point, we can only assume the presence of common properties in a group of IntG genes aimed at achieving an advantage in their expression. A definitive answer to the question, of whether the intracellular production of the HG group of genes decreases with age, can be obtained by studying the changes in the proteome at different periods of life. This will be the main focus of our subsequent work.

## Author contributions

**L. Salnikov** has proposed the theoretical model of aging based on functional genome partition and the role of ratio of the genome two functional parts activity in ontogenesis and aging. He also proposed a way to test the hypothesis using meta-analysis of methylation data. **S. Goldberg** did the statistical analysis of the data. **P. Sukumaran** assisted with data extraction and analysis. **E. Pinsky** organized the work and participated in the data analysis. All authors have made a contribution to prepare the article for publication, participated in the structuring of manuscript and helped with editing. All the authors reviewed, revised and approved the final version of the manuscript.

## Autor Approvales

All authors have seen and approved the manuscript. The manuscript hasn’t been accepted or published elsewhere.

